# Proteomic analysis reveals Utf1 as a neurogenesis-associated new Sumo target

**DOI:** 10.1101/2020.11.17.386557

**Authors:** Juan F. Correa-Vázquez, Francisco Juárez-Vicente, Pablo García-Gutiérrez, Sina V. Barysch, Frauke Melchior, Mario García-Domínguez

**Affiliations:** Andalusian Center for Molecular Biology and Regenerative Medicine-CABIMER CSIC-Universidad de Sevilla-Universidad Pablo de Olavide. Av. Américo Vespucio 24, 41092 Seville, Spain; Zentrum fur Molekulare Biologie Heidelberg (ZMBH), Deutsches Krebsforschungszentrum (DKFZ)-ZMBH Alliance. Im Neuenheimer Feld 282, 69120 Heidelberg, Germany

## Abstract

Post-translational modification by covalent attachment of Sumo regulates numerous processes in vertebrates. Despite demonstrated roles of Sumo in development and function of the nervous system, the identification of key factors displaying a sumoylation-dependent activity during neurogenesis remains elusive. Based on SILAC, we have identified the Sumo proteome of proliferating and neuronal-differentiating cells. More than 300 putative Sumo targets differentially associated with one or the other condition. Among these, Utf1 revealed as a new Sumo target. Gain-of-function experiments demonstrated marked differences between the effects on neurogenesis of wild type and sumoylation mutant versions of selected proteins. While sumoylation of Prox1, Sall4a, Trim24 and Utf1 associated with a positive effect on neurogenesis, sumoylation of Kctd15 associated with a negative effect. Similar results were observed in embryos. Finally, detailed analysis of Utf1 showed sumoylation-dependent control of bivalent genes expression. This effect relies on two mechanisms: sumoylation modulates Utf1 chromatin binding and mediates recruitment of the mRNA-decapping enzyme Dcp1a through a conserved SIM. Altogether, our results indicate that combined sumoylation status of key proteins determine proper progress of neurogenesis.

Sumo/transcription/chromatin/neurogenesis/Utf1

## Introduction

The Small ubiquitin-like modifier (Sumo) is a small polypeptide similar to ubiquitin, capable of covalently attach to other proteins as a post-translational modifier (Flotho & Melchior, 2013). Sumo regulates multiple processes in the eukaryotic cell although it shows a prominent role in transcription repression (Garcia-Dominguez & Reyes, 2009). Among the different Sumo species, Sumo1 normally appears conjugated to proteins, while a large proportion of Sumo2 and Sumo3 (designated as Sumo2/3 due to high homology and similar function) is free and rapidly conjugated to proteins in response to stress conditions (Tempe *et al*, 2008). Sumoylation occurs at lysine residues, often included in the consensus ΨKxE, (Ψ, large hydrophobic residue). For conjugation, proteolytically matured Sumo is activated and transferred to the conjugating enzyme Ubc9, which finally attaches Sumo to target proteins, with the eventual concourse of a Sumo ligase. Sumo maturation and excision from targets is achieved by specific proteases.

Sumoylation is essential in vertebrates, as evidences lethality of the *Ubc9* mutation at early postimplantation stage in the mouse embryo (Nacerddine *et al*, 2005). Sumo government of relevant biological processes makes at present more and more evident the involvement of this modifier in multitude of diseases (Yang *et al*, 2017). In the nervous system, association of Sumo with a variety of neurodegenerative diseases has been shown (Anderson *et al*, 2017; Princz & Tavernarakis, 2020). Besides this, sumoylation plays a fundamental role in the establishment of the synapse, and has revealed as a cytoprotective mechanism in response to severe stress like ischemia (Bernstock *et al*, 2018; Schorova & Martin, 2016). The crucial role Sumo plays during development, including development of the nervous system, is noticeable (Lomeli & Vazquez, 2011). Developmental regulation of the Sumo machinery has been reported for the rodent brain (Hasegawa *et al*, 2014; Loriol *et al*, 2012). However, the role of sumoylation of specific factors in the process of neuronal differentiation has not been deeply investigated. Neurogenesis involves the spatiotemporal deployment of a high number of transcription factors and other proteins subjected to strict regulation (Fujita, 2014; Martynoga *et al*, 2012). We have previously assigned roles in neurogenesis to the Sumo protease Senp7 and to sumoylated Braf35, a component of the LSD1-CoREST histone demethylase complex (Ceballos-Chavez *et al*, 2012; Juarez-Vicente *et al*, 2016), but a detailed landscape of key Sumo targets involved in primary events at early neurogenesis still lacks. To uncover relevant roles of Sumo at the onset of neuronal differentiation, we have determined and compared the Sumo proteomes of proliferating and neuronal-differentiating cells. We have identified a number of proteins whose effect on neurogenesis depends on Sumo attachment. Among these proteins, we have found the transcription factor Utf1, a new Sumo target whose transcription activity depends on sumoylation.

## Results

### Proteomic analysis identifies different subsets of Sumo targets associated with proliferation and neuronal differentiation

To investigate the potential role of sumoylation during early neurogenesis we aimed at identifying and comparing the Sumo proteomes of proliferating and neuronal-differentiating cells. P19 cells were used as a model due to easy manipulation and efficient differentiation, achieved through retinoic acid (RA) treatment or by forced expression of neurogenic factors (Farah *et al*, 2000; McBurney, 1993). For the identification of putative Sumo targets we used the method described in (Barysch *et al*, 2014; Becker *et al*, 2013) to enrich endogenous sumoylated proteins, followed by SILAC-based quantitative mass-spectrometry analysis. We compared cells under proliferation conditions (0 days of RA treatment) with cells treated with RA for 4 days, a stage in which they have formed embryoid bodies and start initial differentiation, mimicking early neurogenesis (Endo *et al*, 2009). Combined cell lysates prepared under denaturing conditions to avoid de-sumoylation by endogenous Sumo were subjected to immunoprecipitation (IP) with Sumo1 or Sumo2/3 antibodies or with normal mouse IgG. We performed two independent experiments. In experiment 1, we used heavy labeling for proliferation conditions and light labeling for differentiation conditions, while in experiment 2 we used reverse labeling (Fig. 1A). Mass spectrometry data (Source Data) permitted the identification of 2240 putative Sumo targets. To define candidates preferentially sumoylated under one or the other condition we established the following criteria: we first considered only proteins giving a SILAC ratio in both experiments (1431 for Sumo1 and 1249 for Sumo2/3), then we asked that ratio in one experiment was inverse to the ratio in the other experiment, and finally, we considered only proteins with a log_2_ (ratio) ≥1 in one experiment and ≤-1 in the other experiment, i.e. a fold-change ≥ 2 in both experiments, with inverse ratios (Fig. 1B). 318 proteins, with variable preference for Sumo1 or Sumo2/3, fitted these criteria (Fig. 1C,D upper panel). Preliminary analysis indicated that most of these proteins were identified on the basis of being better expressed under one or the other condition, in which they are sumoylated. Most of them (74.5%) preferentially associated with differentiation conditions (Fig. 1B,D lower panel). Gene Ontology (GO) analysis indicated similar biological functions for putative Sumo1 and Sumo2/3 targets (Fig. 1E, common categories). Those proteins associated with proliferation grouped to categories related to growth and pluripotency, while those associated with differentiation grouped to categories related to transcription control, chromatin modification, oxidation-reduction metabolism and nervous system development (Fig. 1E).

**Figure 1.**
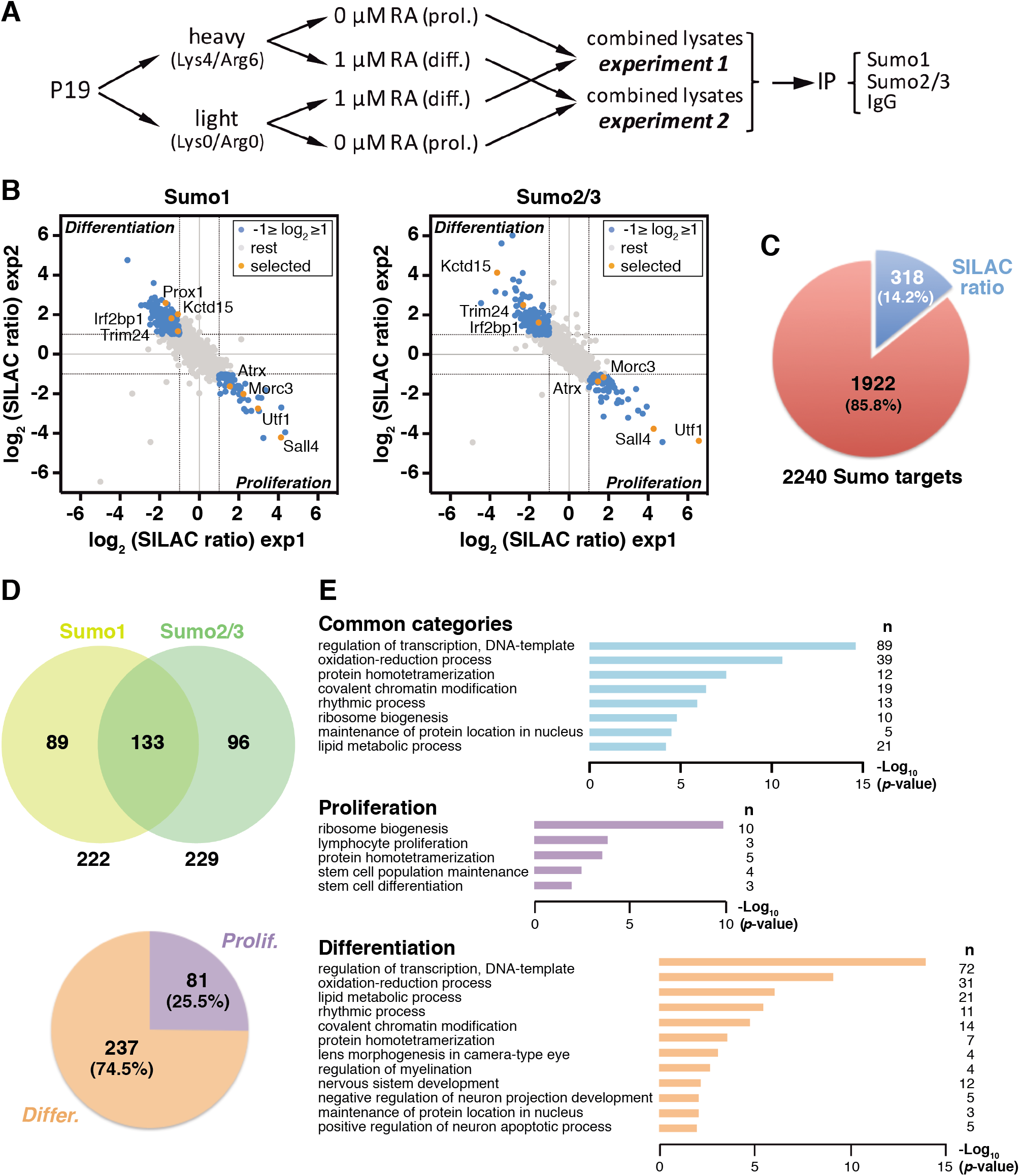
Induction of neuronal differentiation provokes changes in the Sumo proteome. A Schematic representation of the SILAC approach preceding MS analysis. Cells were labeled with heavy or light amino acids under proliferation (prol.) or retinoic acid (RA)-mediated differentiation (diff.), and lysates were combined for immunoprecipitation (IP). B Scatter-plots of Sumo1 and Sumo2/3 putative targets showing a SILAC ratio in both experiments (experiment 1, exp1; experiment 2, exp2). Those with log_2_ (SILAC ratio) ≥1 in one experiment and a log_2_ (SILAC ratio) ≤-1 in the other experiment are in blue dots. Selected proteins for subsequent validation are in orange. The upper-left part of each plot includes proteins more sumoylated under differentiation conditions, while the lower-right part includes proteins more sumoylated under proliferation conditions. C Sector diagram showing the number of proteins indicated in blue in (B), in relation to the total number of Sumo targets identified. D Upper part, Venn diagram showing the number of proteins identified as Sumo1 or Sumo2/3 targets out of 318 proteins. Lower part, sectors diagram showing the proportion of these proteins preferentially sumoylated under proliferation or differentiation conditions. E Gene Ontology analysis (GO) of genes coding for the 318 proteins (common categories), and GO analysis of genes coding for proliferation-or differentiation-associated targets. n, number of genes in each category.

To validate the proteomic approach, we analyzed the sumoylation of selected proteins (Fig. 1B, orange) by using specific antibodies against each protein. Selection was based on observed fold-change and putative functions. Thus, we confirmed the presence of sumoylated Atrx, Morc3, Sall4 and Utf1 in proliferating cells, and of sumoylated Irf2bp1, Kctd15, Prox1 and Trim24 in differentiating cells (Fig. 2AC). Analysis of Brd2 was included as a negative control. We monitored the differentiation process by checking downregulation of the pluripotency marker Oct4 and upregulation of the neuronal marker ßIII-tubulin (Fig. 2B).

**Figure 2.**
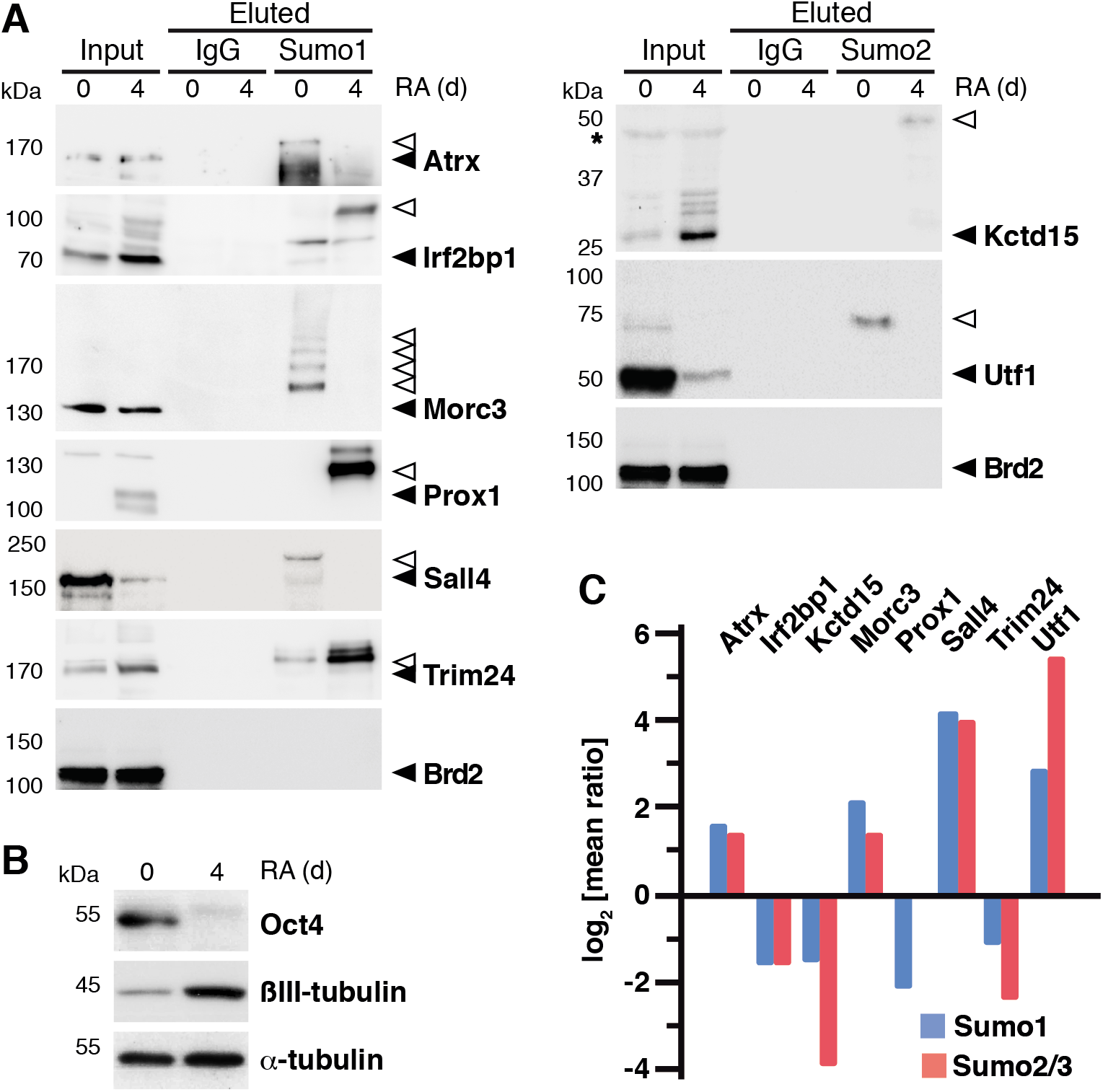
Selected Sumo targets differentially associated with proliferation and differentiation. A Validation of the proteomic results of a selection of proteins by western blot with specific antibodies against the different proteins, after precipitation with Sumo antibodies or with control IgG and peptide elution (100%), under proliferation (0 days (d)) or differentiation conditions (4 d of RA treatment). Some proteins were validated with Sumo1 antibodies and others with Sumo2/3 antibodies. Inputs (1.5%) are also shown. Black arrowheads indicate unmodified proteins while white arrowheads indicate sumoylated products. B Proliferation (0 days) and differentiation conditions (4 days of RA treatment) were checked by analysis of the pluripotency marker Oct4 and the differentiation marker ßIII-tubulin. α-tubulin was determined as a loading marker. C The log2 of the mean value of the ratio in experiment 1 and the inverse ratio in experiment 2 for selected proteins, either for Sumo1 (blue) or for Sumo2/3 (red), is represented.

### Expression of sumoylation mutants of selected proteins alters neuronal differentiation

Among validated proteins, we selected Kctd15, Prox1, Sall4, Trim24 and Utf1 for an initial characterization, based on previously described roles potentially involving them in governing transition from proliferation to differentiation (Dutta & Dawid, 2010; Misra *et al*, 2008; Niederreither *et al*, 1999; Okuda *et al*, 1998; Rao *et al*, 2010). Of these proteins, Kctd15, Prox1, Sall4 and Trim24 were previously shown to be sumoylated (Appikonda *et al*, 2018; Shan *et al*, 2008; Yang *et al*, 2012; Zarelli & Dawid, 2013a), while sumoylation of Utf1 has not been reported so far. Of note, Sall4 sumoylation has been previously shown for Sall4 isoform b (Yang *et al*, 2012). However, in P19 cells we were only able to identify the longest isoform a. Sall4 size in western blot supports this (Fig. 2A). We confirmed in 293T cell lysates sumoylation of expressed Flag- and HA-tagged versions of Sall4a and Utf1, respectively (Fig. EV1). Then, we proceeded to generate Lys to Arg (KR) sumoylation mutants of Utf1. Since software prediction indicated the absence of sumoylation consensus sites, we performed a detailed mutational analysis of the 5 Lys residues present in the mouse Utf1 sequence, which revealed the need to mutate Lys50, 119, and 210, to efficiently prevent sumoylation (Fig. EV1). Sall4a mutations in Lys151, 379 and 846 were according to (Yang *et al*, 2012), and completely abrogated sumoylation (Fig. EV1). We also generated expression constructs of wild type (WT) and KR versions of the other proteins, being mutations according to (Appikonda *et al*, 2018; Hendriks *et al*, 2017; Shan *et al*, 2008; Zarelli & Dawid, 2013a). Next, we checked for similar levels of expressed tagged WT and KR versions of the proteins, for unaltered cell localization and for overexpression levels in relation to the endogenous protein (Fig. EV2).

We decided to study the effects on neurogenesis of KR versus WT versions of selected proteins in gain-of-function experiments. This approach, although involving ectopic expression, helps to have an idea of the impact of sumoylation on activity of Sumo targets. We estimated neurogenesis in P19 cells according to previous reports (Farah *et al*, 2000; Garcia-Gutierrez *et al*, 2014). Thus, we transfected different constructs together with expression vectors for the neurogenic factor NeuroD2 and its co-factor E12. Around 50% of cells expressed the neuronal marker ßIII-tubulin following expression of neurogenic factor, while virtually none of the cells expressed this marker in the absence of this (Fig. 3A). The effect of the different constructs was variable, although effects of WT and KR versions notably differed in all the cases (Fig. 3B, Fig. EV3). In general, KR versions displayed a negative effect, excepting Kctd15-KR, which enhanced neurogenesis. On the other hand, WT Prox1 and Trim24 enhanced neurogenesis, while WT Kctd15 impaired it. WT Sall4a and Utf1 did not affect NeuroD2-promoted differentiation.

**Figure 3.**
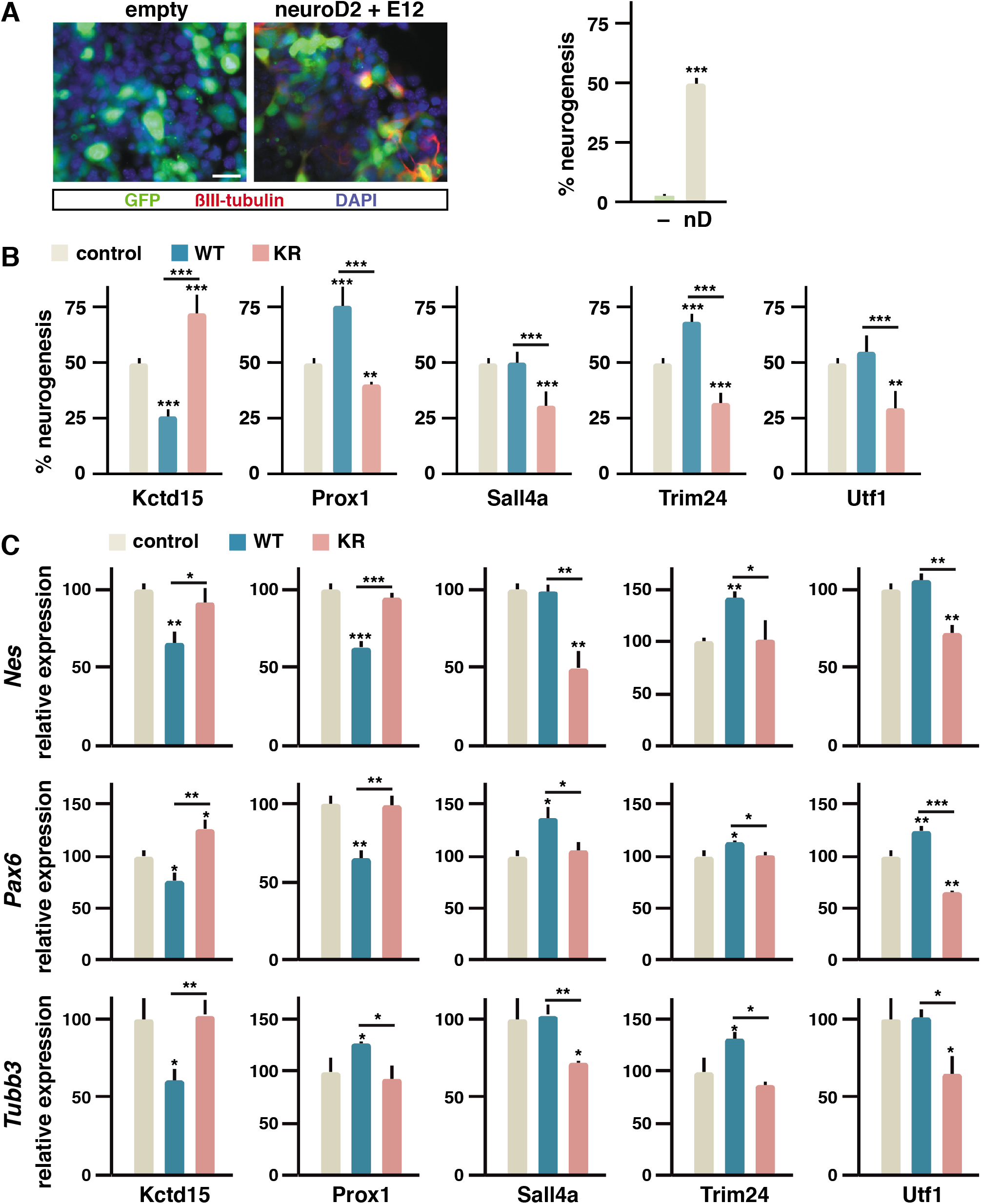
Expression of sumoylation mutants of key Sumo targets alters neurogenesis. A About 50% of neurogenesis is achieved in P19 cells after forced expression of the neurogenic factor NeuroD2 and the E12 co-factor (nD) for 72 hours, in comparison with empty vector transfection (–), as determined in immunofluorescence experiments by the percentage of cells expressing the neuronal marker ßIII-tubulin (red) among transfected cells (green, GFP positive). Nuclei were visualized by DAPI staining (blue). Scale bar 25 μm. B Quantification of the percentage of induced neurogenesis in the presence of expressed wild type (WT) or sumoylation mutant (KR) versions of the indicated proteins. Control corresponds to the percentage of induced neurogenesis in (A). Values are means ± s.d. from counting 140-220 cells in 6 separated fields from 3 independent experiments. C Expression of early (*Nes* and *Pax6*) and late (*Tubb3*) neurogenesis markers was assessed in P19 cells by quantitative PCR after 48 hours of RA-treatment in the absence (control, empty vector) or the presence of expressed WT or KR mutant versions of the indicated proteins. Values are means ± s.d. from 3 independent experiments analyzed in triplicate. Statistical significance in relation to the control is indicated on top of each bar, other comparisons are indicated with a line. Statistical significance was determined by the Student *t*-test. *p*<0.05*, *p*<0.01**, *p*<0.001***.

We also tested the effect of the different constructs on transcription of differentiation markers by quantitative PCR (qPCR) (Fig. 3C). We analyzed expression of early markers *Nes* and *Pax6*, and of later marker *Tubb3.* For convenience we chose to treat the cells with retinoic acid (RA) for 48 hours to promote differentiation. Results were according to those shown in Fig. 3B, with few exceptions: in line with the above results, WT Prox1 promoted *Tubb3* upregulation, but *Nes* and *Pax6* were downregulated, probably due to Prox1-mediated stimulation of neurogenesis negatively affecting early markers at the time of analysis.

### Effect of ectopic Kctd15, Prox1 and Sall4a in the developing neural tube depends on sumoylation

Since Kctd15, Prox1 and Sall4-like proteins have been previously involved in aspects of nervous system development in the vertebrate embryo (Dutta & Dawid, 2010; Misra *et al*, 2008; Young *et al*, 2014), we decided to analyze how sumoylation of these proteins affects the activity they display when expressed in the neural tube. During development, neural progenitors in the neural tube proliferate close to the lumen, the ventricular zone (VZ). They progressively exit the cell cycle and migrate into the pial surface or mantle layer (ML), a ßIII-tubulin positive region where they install to complete differentiation. Migrating early post-mitotic neurons delimit a middle zone between VZ and ML, the subventricular zone (SVZ) (Fig. 4A). We turned to the technique of electroporation of the chick embryo, enabling transfection of expression constructs in neural progenitors of one half of the neural tube. Around 25% of electroporated progenitors in the VZ naturally exit the cell cycle and migrate to the ML (Fig. 4B). However, following ectopic expression of the neurogenic factor Neurogenin 2 (Ngn2), 65% of electroporated cells localized to the ML (Fig. 4B). As in P19 cells, significant differences were observed between the effects of WT and KR versions when co-expressed with Ngn2 (Fig. 4C, Fig. EV4), and similarly, sumoylation of Prox1 and Sall4a associated with neurogenesis stimulation, while sumoylation of Kctd15 associated with neurogenesis impairment. Interestingly, Kctd15-KR promoted neurogenesis in the absence of ectopic Ngn2 (Fig. 4C).

**Figure 4.**
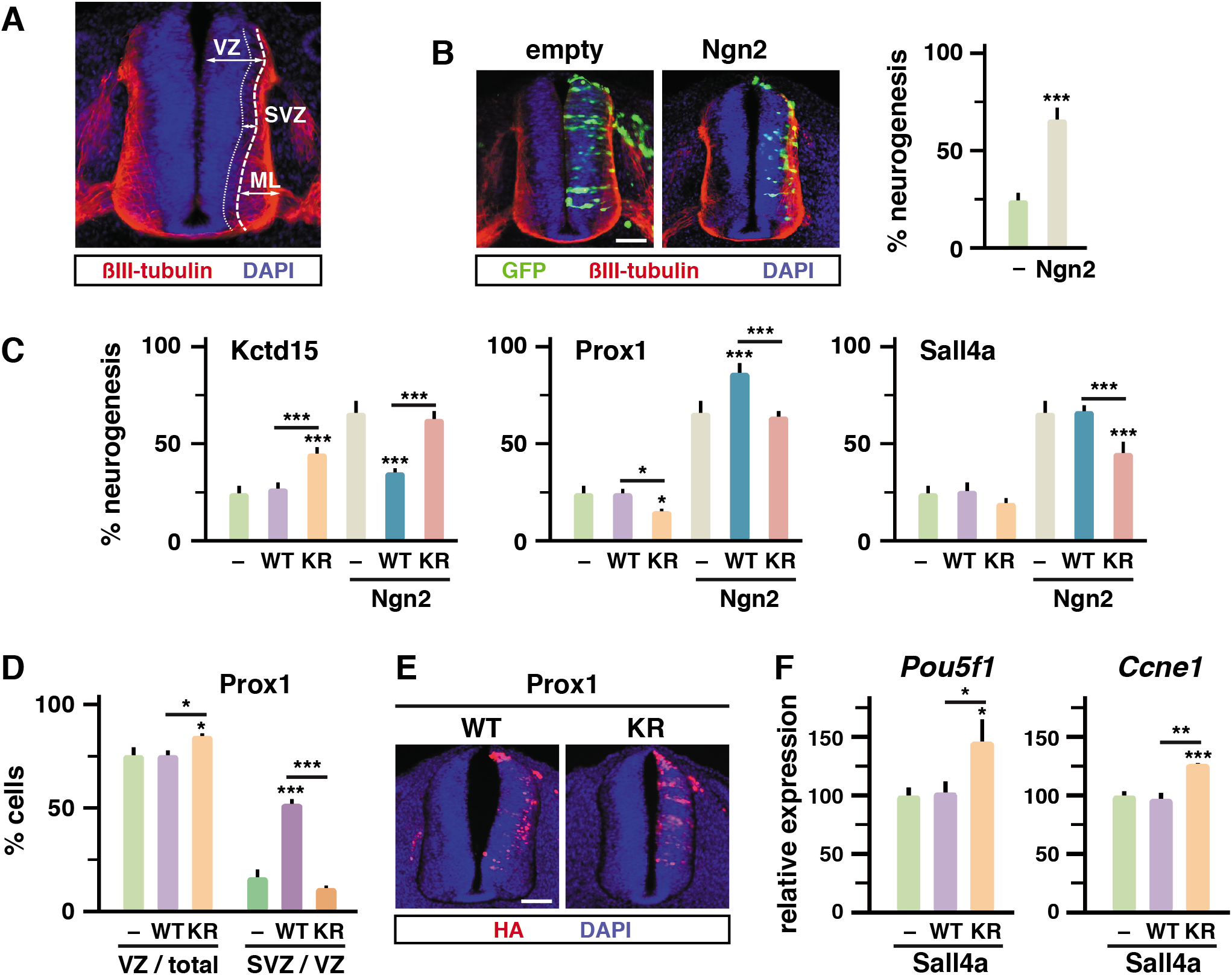
Kctd15, Sall4a and Prox1 display altered effects in the neural tube when mutated for sumoylation. A Section of the developing neural tube indicating the limit between the proliferative ventricular zone (VZ) and the differentiation mantle layer (ML). The sub-ventricular zone (SVZ) of migrating early post-mitotic neurons at the pial surface of the VZ is also indicated. The ML-associated neuronal marker ßIII-tubulin (red) and DAPI-stained nuclei (blue) are shown. B About 65% of neurogenesis is achieved in the developing neural tube after forced expression of the neurogenic factor Neurogenin2 (Ngn2) for 30 hours, in comparison with 25% of naturally occurring neurogenesis when empty vector is electroporated (−). Neurogenesis was determined in immunofluorescence experiments by the percentage of transfected cells (GFP positive, green) located in the ML. C Effect on neurogenesis of WT or KR versions of the indicated proteins, in the absence (empty) or the presence of Ngn2. Controls (−) correspond to the percentages of neurogenesis in the absence (empty) or the presence of Ngn2 as determined in (B). D The percentages of control cells (empty vector electroporation), and of cells expressing WT or KR Prox1 in the absence of Ngn2, localizing to the VZ in relation to the total number of transfected cells, and localizing to the SVZ in relation to the cells localizing to the VZ, were represented. E Localization of expressed HA-tagged WT or KR Prox1 was analyzed by immunofluorescence with anti-HA antibodies (red). Nuclei were visualized by DAPI staining (blue). Scale bars 50 μm. F Expression of *Pou5f1* and *Ccne1* was assessed in P19 cells by quantitative PCR under proliferation conditions in the absence (–, empty vector) or the presence of expressed WT or KR versions of Sall4a. Represented values are means ± s.d. from counting 140-280 cells in 5 sections from 3 independent experiments (B-D) or from 3 independent experiments analyzed in triplicate (F). Statistical significance in relation to the control is indicated on top of each bar, other comparisons are indicated with a line. Statistical significance was determined by ANOVA (*p*<0.0001) followed by Bonferroni’s post-test (95% confidence intervals) (C, D) or by the Student *t*-test. *p*<0.05*, *p*<0.01**, *p*<0.001*** (B, F).

Prox1 has been described to be expressed in the SVZ and to drive neural progenitors out of the cell cycle when ectopically expressed, but not to induce differentiation (Misra *et al*, 2008). In the absence of Ngn2, Prox1-KR provoked a slight decrease of cells in the ML (Fig. 4C) and thereby a slight increase in the VZ (Fig. 4D), while Prox1-WT seemed not to have effects in comparison with control conditions. However, when looking at cell distribution in the VZ, we observed a clear effect of Prox1-WT in driving cells to the SVZ, and remarkably, this effect was abolished when impeding sumoylation (Fig. 4D). We confirmed differential distribution of WT and KR versions of Prox1 by revealing the presence of the expressed constructs (Fig. 4E).

In *Xenopus*, it has been described that Sall4 promotes posterior neural fate by repression of pluripotency *pou5f3* family genes, the closest homologs of mammalian *Pou5f1* (Oct4) (Young *et al*, 2014). Thus, we wondered whether Sall4a might have a similar effect in our model and whether this effect depends on sumoylation. For that, we monitored *Pou5f1* expression in proliferating P19 cells (not forced to differentiate) in control conditions and in the presence of WT and KR versions of Sall4a. Interestingly, expression of the sumoylation mutant led to the increased expression of *Pouf51* and *Ccne1* (cyclin E1) (Fig. 4F), which may explain impaired differentiation by this mutant.

### Sumoylation controls transcriptional activity of Utf1

Utf1 has been described to control pluripotency and differentiation of embryonic stem (ES) and embryonal carcinoma (EC) cells (Lin *et al*, 2012; van den Boom *et al*, 2007). As Utf1 seems restricted to eutherian mammals and it is absent from other vertebrates (Nishimoto *et al*, 2013), we turned to our murine EC model (P19) for detailed analysis. A dual role has been ascribed to Utf1 in the maintenance of the poised state of bivalent genes: on the one hand, its localization to the chromatin limits access of repressor complexes to promoters; on the other hand, Utf1 binds to the mRNA-Decapping enzyme Dcp1a, recruiting it to promoters, for degradation of leakage mRNAs (Jia *et al*, 2012). Thus, we wondered about the impact of Utf1 sumoylation in the control of bivalent gene expression. We first identified a number of bivalent genes regulated by RA-treatment in P19 cells (Fig. EV5). We then analyzed the effect of overexpressing WT and KR versions of Utf1 on the expression levels of bivalent genes in RA-treated P19 cells. As shown in Fig. 5A, in all the cases, marked differences were observed between the effects of WT and KR proteins, excepting on *T* (Brachyury), which was downregulated by RA (Fig. EV5), and on *Rarb*, which is not bound by Utf1 (Jia *et al*, 2012) and was used as a control. Intriguingly, for a set of genes, the WT protein was associated with a positive effect on gene expression in relation to the KR version, while for the other set of genes it was the opposite (Fig. 5A). We next tested some genes from these two different sets, together with *T*, for localization of WT and KR versions of Utf1 to the corresponding promoters. For this, we conducted chromatin immunoprecipitation experiments (ChIP) by precipitating the expressed HA-tagged proteins. Interestingly, we observed that, regardless of the effect on transcription, the sumoylation-deficient mutant bound more strongly to all promoters tested (Fig. 5B). This suggests that sumoylation modulates chromatin association of Utf1.

**Figure 5.**
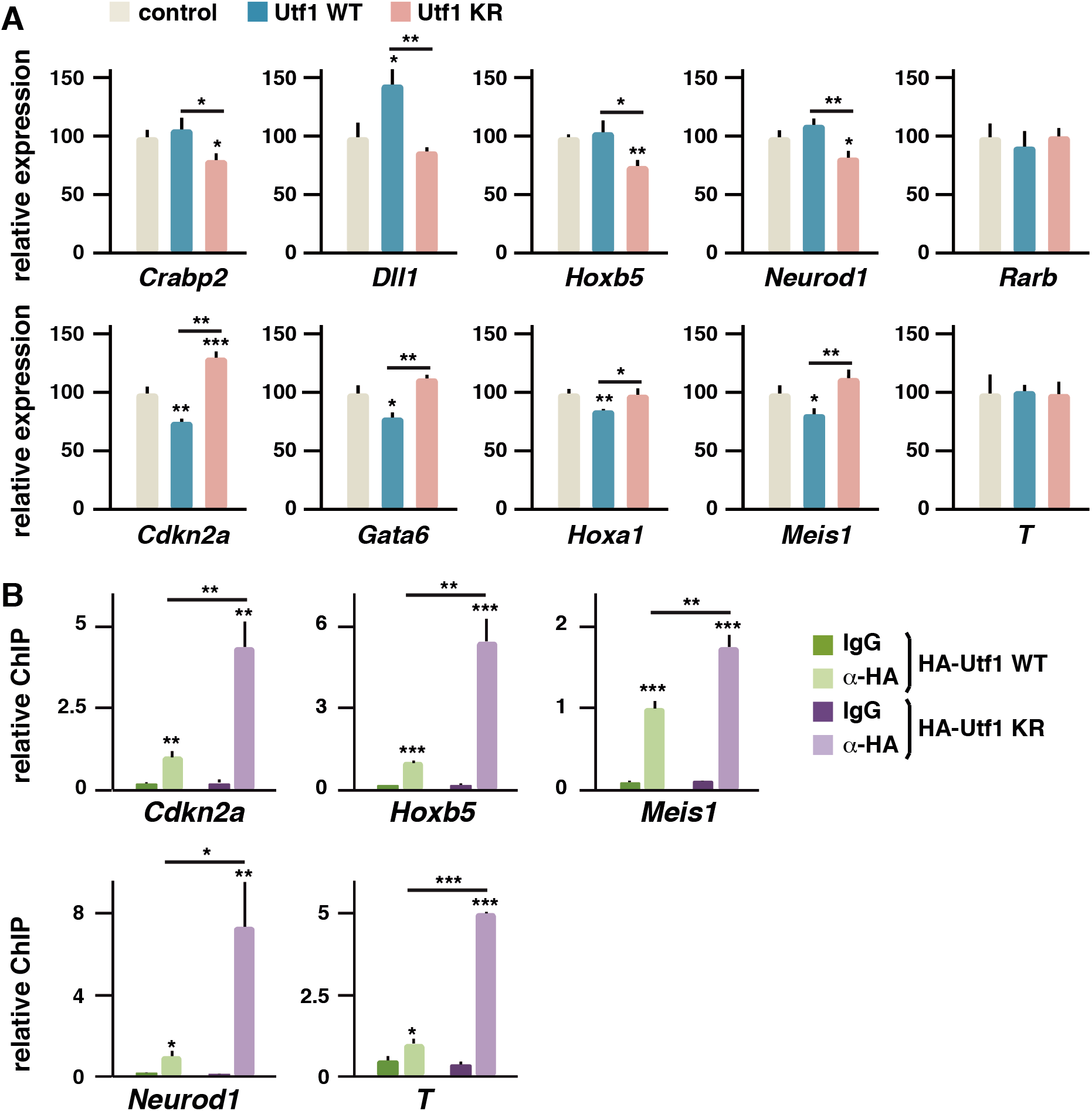
Sumoylation of Utf1 modulates chromatin affinity. A Expression of the indicated bivalent genes was assessed in P19 cells by quantitative PCR after 48 hours of RA-treatment in the absence (control, empty vector) or the presence of expressed WT or KR mutant versions of Utf1. B Localization of expressed HA-tagged WT or KR Utf1 to the indicated promoters was assessed by chromatin immunoprecipitation. IgG controls were established for each expressed construction. Statistical significance in relation to the control (A) or in relation to the corresponding IgG determination (B) is indicated on top of each bar, other comparisons are indicated with a line. Statistical significance was determined by the Student *t*-test. *p*<0.05*, *p*<0.01**, *p*<0.001***.

Finally, we decided to analyze the involvement of Utf1 sumoylation in Dcp1a recruitment. To this purpose, we conducted immunoprecipitation experiments (IP) by precipitating different HA-tagged expressed versions of Utf1 to analyze precipitated endogenous Dcp1a. In addition to WT and KR construct, we prepared a fusion construct of Utf1 with an N-terminal Sumo2 moiety, not cleavable by Sumo proteases. IP demonstrated preferred precipitation of Dcp1a with the Sumo2 fusion (Fig. 6A). Noticeably, analysis of the mouse Dcp1a sequence with the GPS-SUMO 1.0 tool (Zhao *et al*, 2014) revealed the presence of a putative Sumo interacting motif (SIM). This SIM appeared well conserved among vertebrate Dcp1a proteins and displayed scores quite similar to classical Pias1 SIM (Hecker *et al*, 2006) (Fig. 6B). We then tested interaction of the different HA-tagged Utf1 constructs with expressed Flag-tagged Dcp1a, wild type or mutated in the SIM (relevant large hydrophobic residues replaced by alanine). Co-immunoprecipitation experiments revealed interaction of the Sumo2 fusion with wild type, but not with mutated Dcp1a (Fig. 6C). We further tested interaction of the Dcp1a SIM with Sumo2 by a two-hybrid approach. We prepared a bait construct of Sumo2 and prey constructs of the Dcp1a SIM in WT and mutant versions. Growth of yeast in selective medium revealed Sumo interaction with WT SIM but not with the mutant version (Fig. 6D). As a positive control we tested the Pias1 SIM. We finally confirmed this interaction through a pulldown approach (Fig. 6E). Together, these findings suggest that Utf1-Dcp1a interaction can be regulated by SUMOylation.

**Figure 6.**
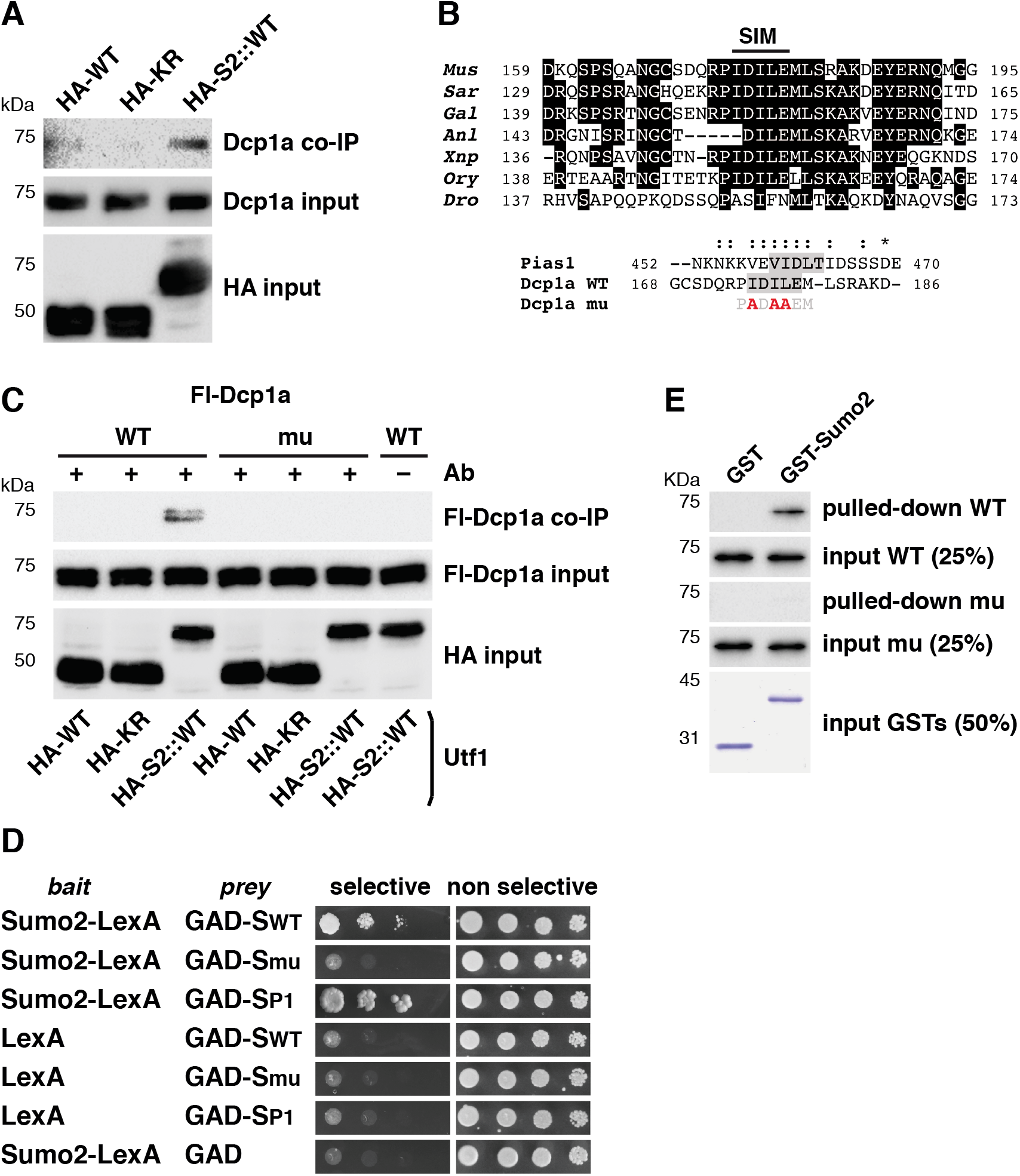
A conserved SIM in Dcp1a mediates interaction with sumoylated Utf1. A Interaction of endogenous Dcp1a with expressed HA-tagged WT, KR or an N-terminal Sumo2 fusion of WT Utf1 (HA-S2::WT) was investigated by co-immunoprecipitation with anti-HA antibodies. Inputs (10%) are also shown. B Alignment of Dcp1a sequences from different species corresponding to a region with a predicted SIM (top, residues conserved in 4 out of the 7 sequences have been boxed in black), and alignment of the mouse Dcp1a WT SIM with the mouse Pias1 SIM (grey shadow), indicating mutations (red) introduced in Dcp1a to disrupt the SIM (Dcp1a mu) (bottom). Similar and identical residues are marked with: and *, respectively. Protein accession numbers: *Mus, Mus musculus* (NP_598522.3); *Sar, Sarcophilus harrisii* (XP_012399256.2); *Gal*, *Gallus gallus* (XP_004944623.1); *Anl, Anolis carolinensis* (XP_008103449.1); *Xnp, Xenopus laevis* (XP_018096411.1); *Ory, Oryzias latipes* (XP_011475154.1); *Dro, Drosophila melanogaster* (NP_611842.1); Pias1, *Mus musculus* (XP_006511365.1). C Interaction of Flag (Fl)-tagged Dcp1a, wild type or mutated in the SIM, with HA-tagged WT, KR or an N-terminal Sumo2 fusion of WT Utf1 (HA-S2::WT) was investigated by co-immunoprecipitation with anti-HA antibodies. Inputs (10%) are also shown. D Two-hybrid assay to probe interaction between Sumo2 fused to the LexA DNA-binding domain (LexA) bait protein and WT (SWT) or mutant (Smu) version of the Dcp1a SIM or the Pias1 SIM (SP1) fused to the GAL4 activation domain (GAD) prey protein. Interaction, as determined by yeast growth, was assessed both in selective and non-selective media. E Flag-tagged wild type (WT) or SIM mutant Dcp1a purified proteins were incubated with GST or a GST-Sumo2 fusion protein. Pulled-down products and the corresponding input were revealed by western blot. GSTs proteins were revealed by Coomassie Blue staining.

## Discussion

To get insights into the role that sumoylation plays during early neurogenesis we have identified and compared the Sumo proteome of proliferating and differentiating cells. Our analysis indicates that a specific pattern of sumoylated proteins differentially define each of these two conditions. Most identified putative Sumo targets consisted in transcription factors and chromatin-associated proteins, which confirms the prevalent transcriptional role of Sumo in regulating relevant cellular processes (Chymkowitch *et al*, 2015). Interestingly, a higher number of sumoylated proteins (ca. 3-fold) associated with differentiation in relation to proliferation, suggesting that Sumo-mediated regulation of a high number of factors is required to initiate such a complex process.

In gain-of-function experiments, we have systematically observed significant differences between WT and KR versions of all proteins on all analyzed markers in the context of neurogenesis. The absence of dramatic effects in some cases may probably be due in part to technical limitations, but also might indicate that sumoylation is involved in fine tuning of protein activities. Thus, the observation of major effects will require global affection of sumoylation or combined sumoylation deficiency of a number of related factors. Indeed, we have previously shown how globally altered sumoylation by overexpression of Sumo1 or Sumo2 in the developing neural tube has drastic consequences on neurogenesis (Juarez-Vicente *et al*, 2016). Results from forced expression of sumoylation mutants have been essential in raising our conclusions. In some cases, these mutants lacked the effect displayed by the corresponding WT version, indicating that effects depend on sumoylation. However, for certain proteins, the KR version behaved as a dominant negative, provoking a clear effect in comparison with control conditions, which probably indicates a competition with the endogenous sumoylable protein.

In general, we have been able to associate sumoylation of selected proteins with progression of neurogenesis. Some of the best SUMO targets that we have identified here are proteins that are upregulated under differentiation conditions and which contribute to neurogenesis. Our finding that they are strong SUMO targets indicates that regulation by SUMO could be an important concept in the temporal progression of neurogenesis. Both Trim24 and Prox1 were more expressed under differentiation conditions, correlating their sumoylation with a positive role in neurogenesis. According to a previous report (Misra *et al*, 2008) we observe Prox1-WT-mediated driving of cells to the SVZ in the absence of Ngn2. In addition, we have observed a positive effect in neurogenesis in the presence of Ngn2. Remarkably, both effects depend on sumoylation. Sall4a and Utf1, which were better expressed under proliferation conditions, also demonstrate to have a positive role in neurogenesis when sumoylated. Interestingly, we have detected *Sall4a* but not *Sall4b* transcript in P19 cells. While Sall4b has been more related to pluripotency, Sall4a has been related to differentiation and patterning, and more precisely to specific embryonic layers, in particular ectoderm (Rao *et al*, 2010).

Sumo attachment to Utf1 seems to participate in at least two functional aspects: a) modulating affinity for chromatin, and b) facilitating binding to Dcp1a. Thus, Utf1 adds to the list of transcription factors, sumoylation of which regulates its transcription activity. The effects of WT and KR proteins on expression of bivalent genes were markedly different. The negative effect of KR Utf1 in neurogenesis correlates with the previously reported role of WT protein in stem cells differentiation and in transcriptional control of bivalent genes (Jia *et al*, 2012; van den Boom *et al*, 2007). Sumo, by avoiding strong localization of Utf1 to regulated promoters, should facilitate partial access of repressors, but also would assure Dcp1a recruitment to promoters for decapping activity on leakage mRNAs in order to maintain repression of bivalent genes, but in a poised state for rapid and effective activation upon induction of differentiation. Indeed, enhanced localization of the KR mutant to the chromatin might interfere with localization of neurogenesis-related factors and contribute to explain altered gene expression. Intriguingly, while a set of analyzed genes was negatively affected by the KR, the other set was negatively affected by the WT. However, this correlates with previously reported effect of *Utf1* knockout on gene expression. Thus, those genes negatively affected by Utf1-WT (*Cdkn2a, Gata6, Hoxa1, Meis1*) are genes upregulated upon *Utf1* knockout, while most of those negatively affected by Utf1-KR (*Crabp2, Dll1, Neurod1*) are downregulated upon knockout (Jia *et al*, 2012). Dcp1a interaction with Utf1 seems to be mediated by a SIM sequence conserved in vertebrate proteins. Since Utf1 evolutionarily appeared in eutherian mammals (Nishimoto *et al*, 2013), this raises the question of this SIM present in Dcp1a from non-eutherian mammals and other vertebrates. A possibility is that this SIM is used for interaction with other sumoylated proteins but also for interaction with an ancestor of eutherian Utf1 in other vertebrates.

Finally, Kctd15, preferentially expressed and sumoylated under differentiation conditions, appeared to be involved in delaying neurogenesis. This suggests the involvement of Kctd15 sumoylation in controlling proper timing for neurogenesis progression, that otherwise might result in premature aberrant differentiation. Of note, we recently described another mechanism, also involved in delaying neurogenesis to avoid abrupt and abnormal differentiation (Luna-Pelaez & Garcia-Dominguez, 2018). In the embryo, non-sumoylated Kctd15 has been involved in inhibiting neural crest formation (Dutta & Dawid, 2010; Zarelli & Dawid, 2013a; Zarelli & Dawid, 2013b), ascribing then multiple developmental roles to Kctd15 sumoylation. Interestingly, Kctd15-KR showed neurogenic activity in the neural tube, since it promoted neurogenesis in the absence of Ngn2, pointing to a dominant negative effect of the mutant protein, and indicating the probable involvement of progenitors-associated endogenous protein in preventing differentiation.

We cannot exclude additional modifications, others than sumoylation, targeting the same Lys residues studied in this work, but several lines of evidence strongly support the participation of sumoylation in neurogenesis. Future work in this matter will help to define the exact molecular mechanisms underlying the consequences of Sumo attachment to each specific factor and to groups of related factors.

## Material and methods

### Endogenous Sumo immunoprecipitation, SILAC analysis and mass spectrometry

Endogenous Sumo immunoprecipitation was performed as described in great detail (Becker *et al*, 2013). Hybridoma producing monoclonal antibodies SUMO1 (clone 21C7) and SUMO2/3 (Clone 8A2), developed by Dr. Mike Matunis (Matunis *et al*, 1996; Zhang *et al*, 2008), were obtained from the Developmental Studies Hybridoma Bank developed under the auspices of the NICHD and maintained by The University of Iowa, Department of Biology, Iowa City, IA 52242. Cultivation and antibody purification were done as described in (Barysch *et al*, 2014; Becker *et al*, 2013). For the quantification of the Sumo-proteome upon treatment/Mass spectometry, P19 cells were grown for 6-7 doublings (from 30 cm^2^ to 3600 cm^2^ culture dish surface, i.e. 24 15 wells in the end) in SILAC DMEM medium containing dialyzed FBS (dialyzed 3x against PBS through a 6-8000 MWCO bag), 2 mM L-glutamine and 146 μg/ml lysine and 86 μg/ml arginine (conditions under which no undesired amino acid conversion, such as proline to arginine, could be detected). One set of 24 plates contained “light” lysine and arginine, the other set of 24 plates contained D4-lysine and 13C-arginine. 16-18 hours before collecting the cells, they were serum starved in SILAC DMEM medium containing only glutamine and the respective type of lysine and arginine. One set of cells was treated with all *trans* retinoic acid (RA) at 1 μM for 4 days; the other set was treated with vehicle. For large-scale SUMO-IPs, the cells were lysed in 350 μl 2x lysis buffer per plate and lysates from all 48 plates were combined. SUMO-IPs were performed as stated above and TCA precipitated eluates were loaded onto NuPAGE® Novex® Bis-Tris Mini Gels (4-20%) and stained with Coomassie Brilliant Blue. In-gel digestion was performed as described (Shevchenko *et al*, 2006) with minor modifications. Subsequently, peptide extraction and mass spectrometry were performed similar to (Barysch *et al*, 2014; Becker *et al*, 2013). Two independent experiments were performed. In experiment 1, heavy labeling was used for proliferation conditions and light labeling for differentiation conditions, while in experiment 2 reverse labeling was used. We considered only proteins giving a SILAC ratio in both experiments, being the ratio in one experiment inverse to the ratio in the other experiment, both with a log_2_ value ≥ |1|.

### Cell culture, transfection, immunofluorescence and embryo electroporation

Human 293T and mouse P19 cells were cultured in Dulbecco’s modified Eagle’s medium (Sigma-Aldrich, St. Louis, MO, USA) supplemented with 10% fetal bovine serum (Sigma-Aldrich) and alpha-modified Eagle’s medium (Hyclone, Logan, UT, USA) supplemented with 7.5% calf (Hyclone) and 2.5% fetal bovine sera, respectively and 10 ml/l of a antibiotic solution with Penicillin (100 U/ml) and Streptomycin (10 mg/ml) (Sigma-Aldrich). P19 cells were directly obtained from ATCC (Teddington, Middlesex, UK). Transfections were performed with Lipofectamine 2000 (Invitrogen, Life Technologies, Paisley, UK) 36 h before harvesting the cells. All transfection constructs, except the GFP expression vector pEGFP_C2 (BD Biosciences Clontech, San Jose, CA, USA), were derived from vector pAdRSV-Sp (Giudicelli *et al*, 2003), in which different cDNAs were cloned with Flag or HA tags. Mouse cDNAs were cloned by standard PCR procedures from a cDNA preparation from P19 cells-derived total RNA. PCR for cloning was performed with the Q5 polymerase (New England Biolabs, Ipswich, MA, USA). Electroporation, preparation of embryos and immunofluorescence on neural tube sections or P19 cells were performed as previously described (Farah *et al*, 2000; Garcia-Dominguez *et al*, 2003).

### Western blot, immunoprecipitation, pull-down and yeast two-hybrid

Cell extracts were prepared in SDS-containing denaturing buffer (Becker *et al*, 2013), sonicated, immunobloted with the Trans-Blot Turbo transfer system (Biorad, Hercules, CA, USA), and analysed using the Clarity Western ECL Substrate (Biorad) in the Chemidoc XRS Imaging system (Biorad). Brd2 antibody was described in (Garcia-Gutierrez *et al*, 2012). Commercial primary antibodies (1:1000) were as follows: Atrx (sc-15408, Santa Cruz Btg., Dallas, TX, USA), Irf2bp1 (18847-1-AP, Proteintech, Manchester, UK), Morc3 (sc-83730, Santa Cruz Btg.), Prox1 (07-537, Upstate, Merck Millipore, Burlington, MA, USA), Sall4 (ab29112, Abcam, Cambridge, UK), Trim24 (14208-1-AP, Proteintech), Oct4 (H134, sc-9081, Santa Cruz Btg.), ßIII-tubulin (ab18207, Abcam), α-tubulin (DM1A, T9026, Sigma-Aldrich), Kctd15 (PA5-25862, Thermo Fisher Sci., Waltham, MA, USA), Utf1 (ab24273, Abcam), Dcp1a (WH0055802M6, Sigma-Aldrich), HA (H6908, Sigma-Aldrich), Flag (M2, F1804, Sigma-Aldrich). HRP-conjugated antibodies were from Sigma-Aldrich (1:10 000). Yeast two hybrid assay was performed using the DUALhybrid Kit system (Biotech, Schlieren, Zurich, Switzerland), using the pLexA-N bait vector and the pGAD-HA prey vector. Production of proteins was performed in *E. coli* DH5α strain and purification of GST fusion proteins was achieved by incubation with Gluthatione Sepharose 4B matrix (GE Healthcare, Buckinghamshire, UK). Sumo proteins were kept bound to matrix, while Dcp1a was eluted by excision of the GST moiety trough PreScission protease (GE Healthcare) incubation. Pull-down experiments were conducted as previously described (Garcia-Dominguez *et al*, 2008).

### RNA extraction, quantitative PCR and chromatin immunoprecipitation

Total RNA was prepared using RNeasy mini kit (Qiagen, Austin, TX, USA), using the RNase-Free DNase Set (Qiagen) on column following instructions from manufacturers and cDNAs were synthesized using the iSCRIPT cDNA Synthesis Kit (Biorad). Quantitative-PCR (qPCR) was performed with Power SYBR Green on the 7500 Fast Real-Time PCR System (Applied Biosystems, Carlsbad, CA, USA). Chromatin immunoprecipitation was performed as previously described (Garcia-Gutierrez *et al*, 2014). Relative quantities of gene expression levels were normalized to the expression of the housekeeping gene *Rplp0.* Primers are indicated in Appendix Table S1.

### Statistical analysis

One-way ANOVA (*p*<0.05) or *t*-test were applied for statistical analysis as indicated. *p*<0.05*, *p*<0.01**, *p*<0.001***.

## Acknowledgements

This work was supported by grants BFU2015-64721-P and PGC2018-094232-B-I00 from Ministry of Science, Innovation and Universities (MICIU), Spain, to M. G.-D., and also received funding from the Deutsche Forschungsgemeinschaft (DFG, German Research Foundation) - Project Number 278001972 - TRR 186. J.F. C.-V. and F. J.-V. were the recipients of FPI (MICIU, BES-2016-076500) and JAE PhD (CSIC) fellowships, respectively. We acknowledge the ZMBH Core Facility for Mass Spectrometry & Proteomics. We thank Dr. Annette Flotho for help with data analysis. We also acknowledge LM Buch for preliminary results on Kctd15.

## Author Contributions

JF C-V, validated positives, performed expression analysis, ChIPs, IPs and electroporation. F J-V, performed the proteomic approach, validated positives, performed electroporation. P G-G, performed protein purification and pull-down, helped with cell culture and IPs. SV B, helped in proteomic analysis and validations, analyzed data, supervised experiments and edited the manuscript. F M, designed and supervised experiments and contributed to build the manuscript. M G-D, conceived the project, designed and supervised experiments, contributed to some experiments, analyzed data, wrote the manuscript and assembled the figures.

## Expanded View Figure legends

**Figure EV1.**
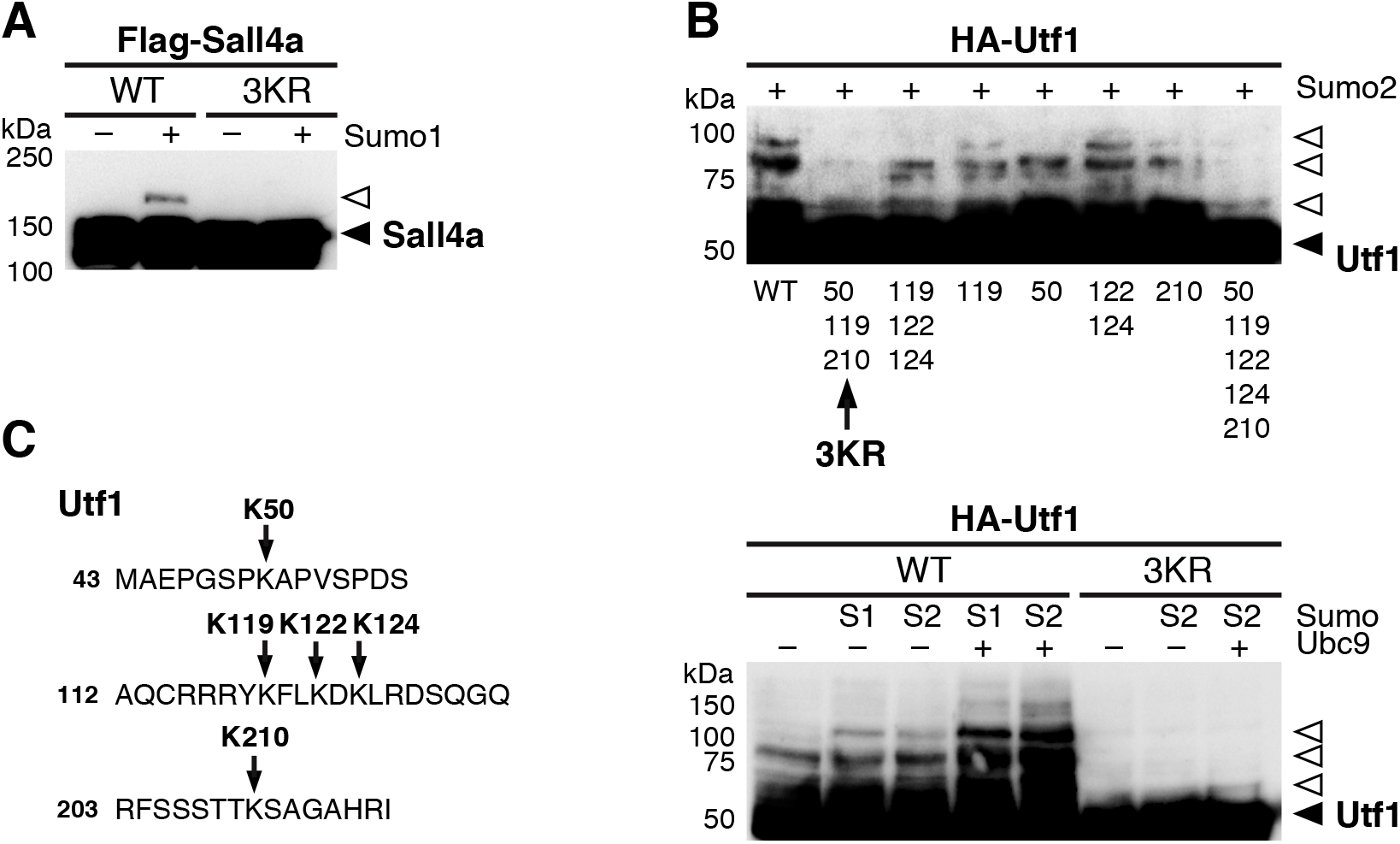
Sall4 and Utf1 sumoylation in cells. A, B Sumoylation of expressed Flag-tagged Sall4a (A) and HA-tagged Utf1 (B) was tested in 293T cells through western blot with anti-Flag and anti-HA antibodies, respectively, in the absence (−) or the presence of expressed Ubc9 (+) and the indicated Sumo species (S1, Sumo1; S2, Sumo2). Absence of sumoylation of expressed sumoylation mutants was also tested (Flag-Sall4a 3KR and HA-Utf1 3KR). Black arrowheads indicate unmodified proteins while white arrowheads indicate sumoylated products. 20 μg of total protein were loaded per lane. C Position of the 5 Lys residues present in the mouse Utf1 sequence and mutated for the analysis in (B), are indicated.

**Figure EV2.**
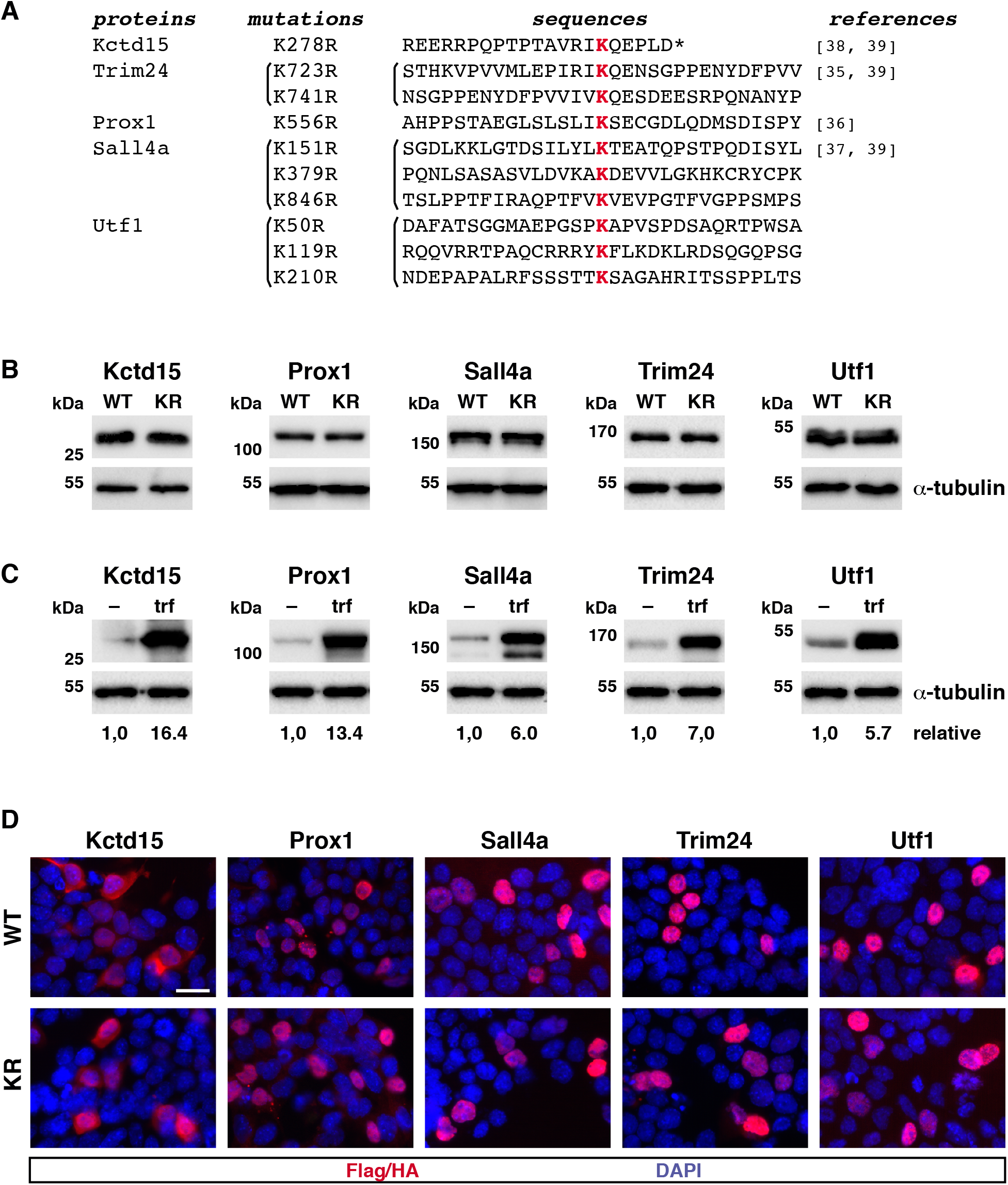
Expression and cellular localization of WT and KR sumoylation mutants of Kctd15, Prox1, Sall4a, Trim24 and Utf1 in P19 cells. A Mutated K (red) and surrounding sequences of selected proteins and references for previously described mutations. B Level of expression of expressed wild type (WT) or sumoylation mutant (KR) versions of Flag-tagged Kctd15, Sall4a and Trim24, or HA-tagged Prox1 and Utf1 proteins were determined in P19 cells by western blot. C Fold-overexpression was determined for each WT expression construct in western blot experiments of transfected (trf) and non-transfected (–) cells with specific antibodies against each protein. α-tubulin was determined as a loading marker. 20 μg of total protein were loaded per lane. D Cellular localization of Flag-or HA-tagged proteins analyzed by immunofluorescence with anti-Flag or anti-HA antibodies (red), respectively. Nuclei were visualized by DAPI staining (blue). Scale bar 25 μm.

**Figure EV3.**
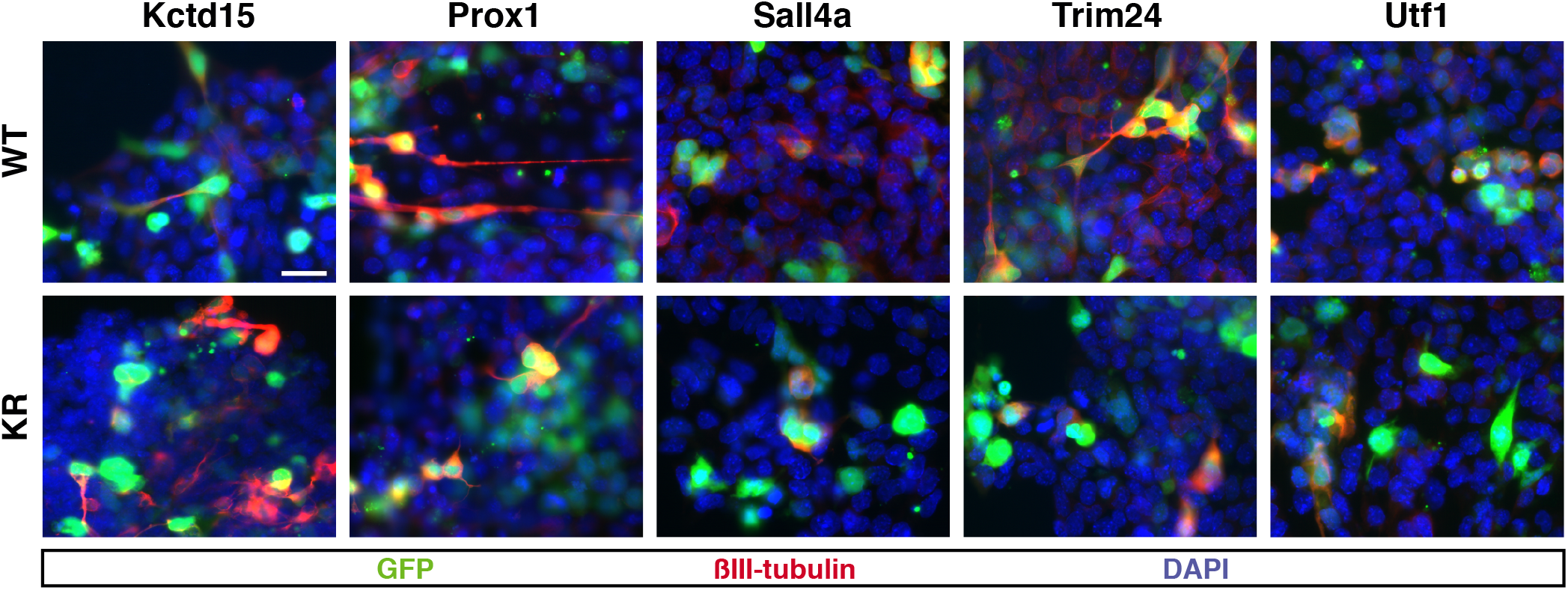
Neurogenesis analysis in P19 cells. P19 cells were transfected with expression cosntructs for NeuroD2 and the E12 co-factor together expression constructs for WT or KR versions of the indicated proteins. Neurogenesis was evaluated 72 hours later by revealing the neuronal marker βIII-tubulin (red). Transfected cells were visualized by expression of a GFP reporter. Nuclei were visualized by DAPI staining (blue). Scale bar 25 μm.

**Figure EV4.**
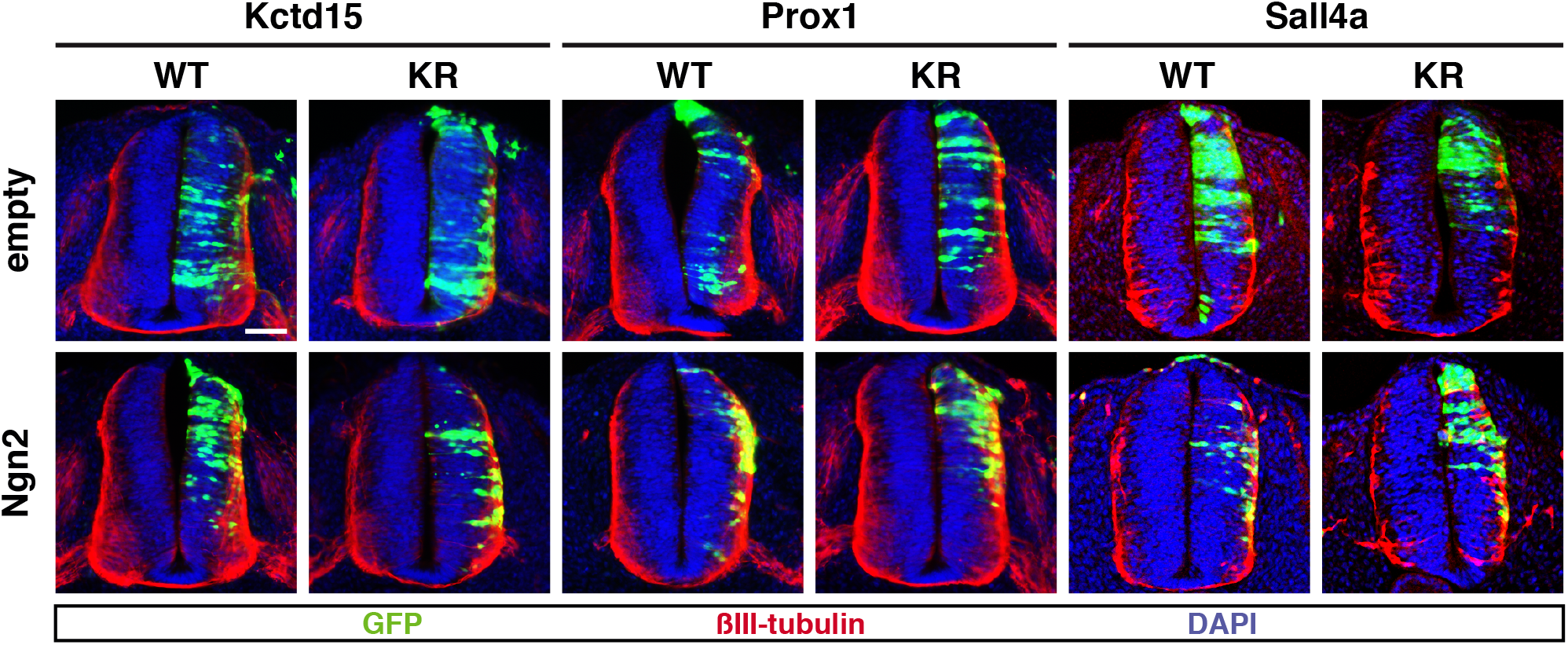
Neurogenesis analysis in embryos. Neurogenesis in the developing neural tube was analyzed on embryos electroporated for 30 h with expression constructs for WT or KR versions of the indicated proteins, in the absence (empty) or the presence of a expression construct for the neurogenic factor Neurogenin2 (Ngn2). Mantle layer was visualized by revealing the marker βIII-tubulin (red). Nuclei were marked by DAPI-staining (blue). Scale bar 50 μm.

**Figure EV5.**
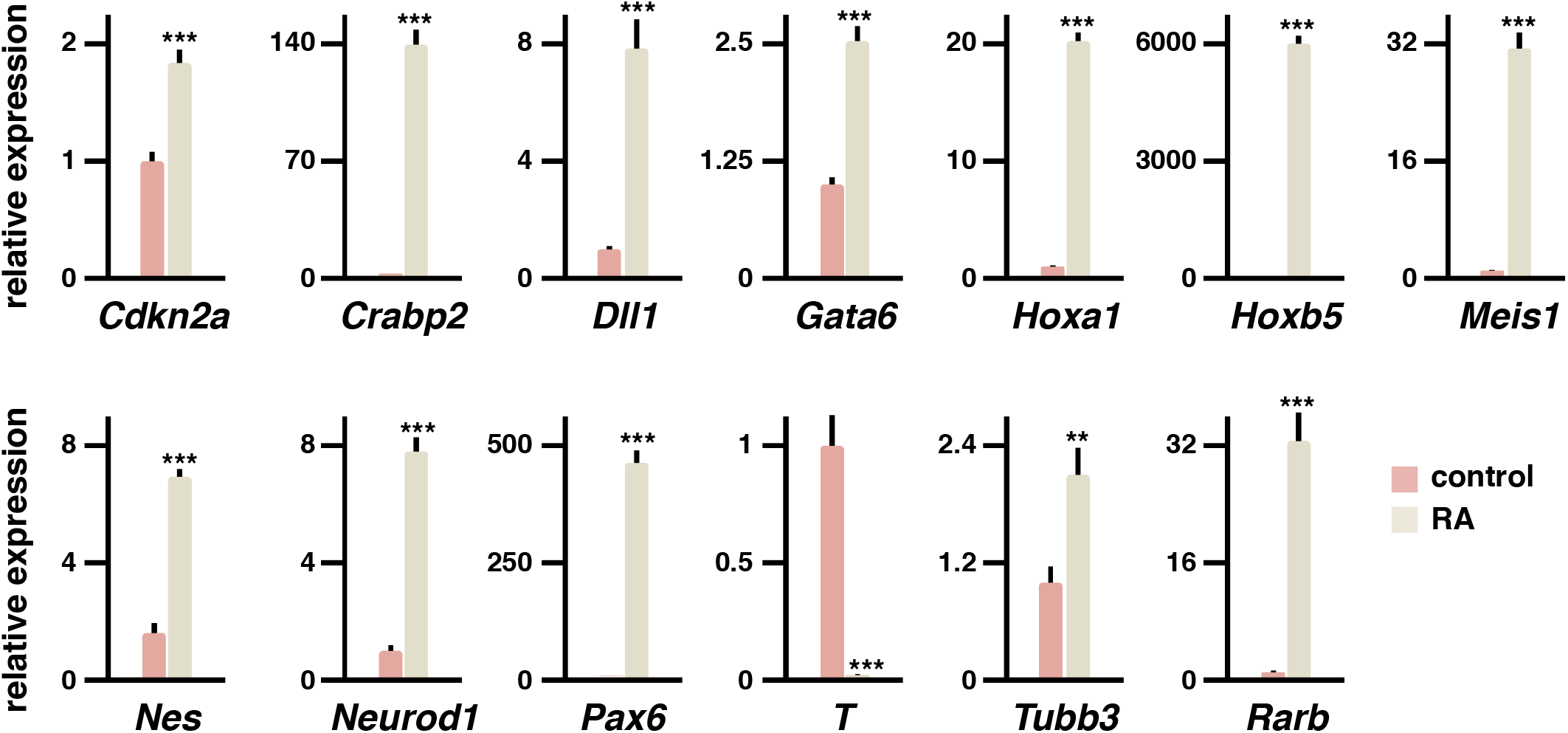
Retinoic acid regulation of bivalent gene expression in P19 cells. Levels of expression of the indicated bivalent genes were assessed in P19 cells by quantitative PCR under control proliferation conditions or after 48 hours of RA-treatment. Values are means ± s.d. from 3 independent experiments analyzed in triplicate. Statistical significance in relation to the control is indicated on top of each bar. Statistical significance was determined by the Student *t*-test. *p*<0.05*, *p*<0.01**, *p*<0.001***.

**Appendix Table S1.**
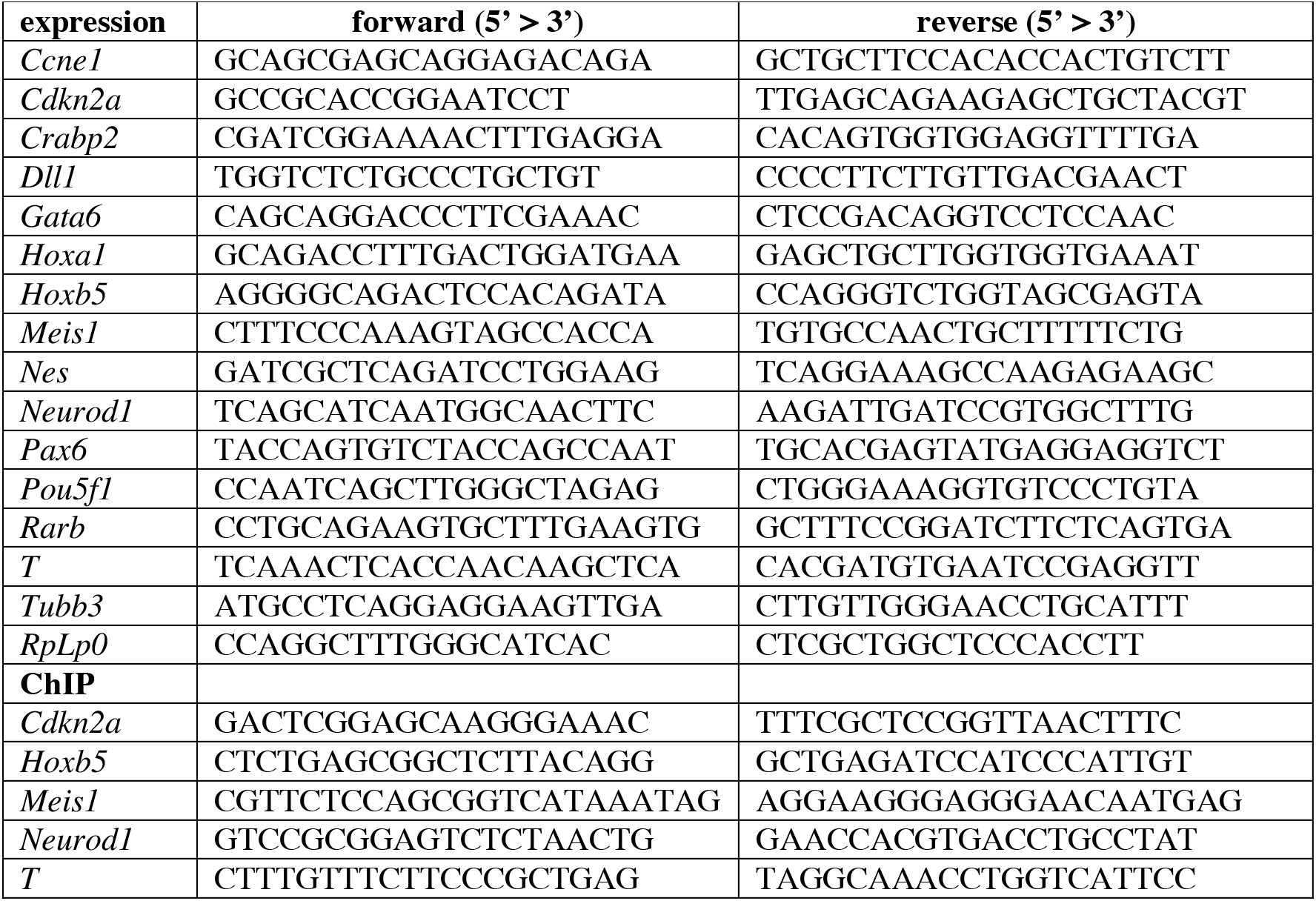
Primers used for expression and ChIP analyses

## References

Anderson DB, Zanella CA, Henley JM, Cimarosti H (2017) Sumoylation: Implications for Neurodegenerative Diseases. Adv Exp Med Biol 963: 261–281

Appikonda S, Thakkar KN, Shah PK, Dent SYR, Andersen JN, Barton MC (2018) Cross-talk between chromatin acetylation and SUMOylation of tripartite motif-containing protein 24 (TRIM24) impacts cell adhesion. J Biol Chem 293: 7476–7485

Barysch SV, Dittner C, Flotho A, Becker J, Melchior F (2014) Identification and analysis of endogenous SUMO1 and SUMO2/3 targets in mammalian cells and tissues using monoclonal antibodies. Nat Protoc 9: 896–909

Becker J, Barysch SV, Karaca S, Dittner C, Hsiao HH, Berriel Diaz M, Herzig S, Urlaub H, Melchior F (2013) Detecting endogenous SUMO targets in mammalian cells and tissues. Nat Struct Mol Biol 20: 525–531

Bernstock JD, Yang W, Ye DG, Shen Y, Pluchino S, Lee YJ, Hallenbeck JM, Paschen W (2018) SUMOylation in brain ischemia: Patterns, targets, and translational implications. J Cereb Blood Flow Metab 38: 5–16

Ceballos-Chavez M, Rivero S, Garcia-Gutierrez P, Rodriguez-Paredes M, Garcia-Dominguez M, Bhattacharya S, Reyes JC (2012) Control of neuronal differentiation by sumoylation of BRAF35, a subunit of the LSD1-CoREST histone demethylase complex. Proc Natl Acad Sci USA 109: 8085–8090

Chymkowitch P, Nguea PA, Enserink JM (2015) SUMO-regulated transcription: challenging the dogma. Bioessays 37: 1095–1105

Dutta S, Dawid IB (2010) Kctd15 inhibits neural crest formation by attenuating Wnt/beta-catenin signaling output. Development 137: 3013–3018

Endo M, Antonyak MA, Cerione RA (2009) Cdc42-mTOR signaling pathway controls Hes5 and Pax6 expression in retinoic acid-dependent neural differentiation. J Biol Chem 284: 5107–5118

Farah MH, Olson JM, Sucic HB, Hume RI, Tapscott SJ, Turner DL (2000) Generation of neurons by transient expression of neural bHLH proteins in mammalian cells. Development 127: 693–702

Flotho A, Melchior F (2013) Sumoylation: a regulatory protein modification in health and disease. Annu Rev Biochem 82: 357–385

Fujita S (2014) 50 years of research on the phenomena and epigenetic mechanism of neurogenesis. Neurosci Res 86: 3–13

Garcia-Dominguez M, March-Diaz R, Reyes JC (2008) The PHD domain of plant PIAS proteins mediates sumoylation of bromodomain GTE proteins. J Biol Chem 283: 21469–21477

Garcia-Dominguez M, Poquet C, Garel S, Charnay P (2003) Ebf gene function is required for coupling neuronal differentiation and cell cycle exit. Development 130: 6013–6025

Garcia-Dominguez M, Reyes JC (2009) SUMO association with repressor complexes, emerging routes for transcriptional control. Biochim Biophys Acta 1789: 451–459

Garcia-Gutierrez P, Juarez-Vicente F, Wolgemuth DJ, Garcia-Dominguez M (2014) Pleiotrophin antagonizes Brd2 during neuronal differentiation. J Cell Sci 127: 2554–2564

Garcia-Gutierrez P, Mundi M, Garcia-Dominguez M (2012) Association of bromodomain BET proteins with chromatin requires dimerization through the conserved motif B. J Cell Sci 125: 3671–3680

Giudicelli F, Gilardi-Hebenstreit P, Mechta-Grigoriou F, Poquet C, Charnay P (2003) Novel activities of Mafb underlie its dual role in hindbrain segmentation and regional specification. Dev Biol 253: 150–162

Hasegawa Y, Yoshida D, Nakamura Y, Sakakibara S (2014) Spatiotemporal distribution of SUMOylation components during mouse brain development. J Comp Neurol 522: 3020–3036

Hecker CM, Rabiller M, Haglund K, Bayer P, Dikic I (2006) Specification of SUMO1- and SUMO2-interacting motifs. J Biol Chem 281: 16117–16127

Hendriks IA, Lyon D, Young C, Jensen LJ, Vertegaal AC, Nielsen ML (2017) Site-specific mapping of the human SUMO proteome reveals co-modification with phosphorylation. Nat Struct Mol Biol 24: 325–336

Jia J, Zheng X, Hu G, Cui K, Zhang J, Zhang A, Jiang H, Lu B, Yates J, 3rd, Liu C, et al (2012) Regulation of pluripotency and self-renewal of ESCs through epigenetic-threshold modulation and mRNA pruning. Cell 151: 576–589

Juarez-Vicente F, Luna-Pelaez N, Garcia-Dominguez M (2016) The Sumo protease Senp7 is required for proper neuronal differentiation. Biochim Biophys Acta 1863: 1490–1498

Lin CH, Yang CH, Chen YR (2012) UTF1 deficiency promotes retinoic acid-induced neuronal differentiation in P19 embryonal carcinoma cells. Int J Biochem Cell Biol 44: 350–357

Lomeli H, Vazquez M (2011) Emerging roles of the SUMO pathway in development. Cell Mol Life Sci 68: 4045–4064

Loriol C, Parisot J, Poupon G, Gwizdek C, Martin S (2012) Developmental regulation and spatiotemporal redistribution of the sumoylation machinery in the rat central nervous system. PLoS One 7: e33757

Luna-Pelaez N, Garcia-Dominguez M (2018) Lyar-Mediated Recruitment of Brd2 to the Chromatin Attenuates Nanog Downregulation Following Induction of Differentiation. J Mol Biol 430: 1084–1097

Martynoga B, Drechsel D, Guillemot F (2012) Molecular control of neurogenesis: a view from the mammalian cerebral cortex. Cold Spring Harb Perspect Biol 4

Matunis MJ, Coutavas E, Blobel G (1996) A novel ubiquitin-like modification modulates the partitioning of the Ran-GTPase-activating protein RanGAP1 between the cytosol and the nuclear pore complex. J Cell Biol 135: 1457–1470

McBurney MW (1993) P19 embryonal carcinoma cells. Int J Dev Biol 37: 135–140

Misra K, Gui H, Matise MP (2008) Prox1 regulates a transitory state for interneuron neurogenesis in the spinal cord. Dev Dyn 237: 393–402

Nacerddine K, Lehembre F, Bhaumik M, Artus J, Cohen-Tannoudji M, Babinet C, Pandolfi PP, Dejean A (2005) The SUMO pathway is essential for nuclear integrity and chromosome segregation in mice. Dev Cell 9: 769–779

Niederreither K, Remboutsika E, Gansmuller A, Losson R, Dolle P (1999) Expression of the transcriptional intermediary factor TIF1alpha during mouse development and in the reproductive organs. Mech Dev 88: 111–117

Nishimoto M, Katano M, Yamagishi T, Hishida T, Kamon M, Suzuki A, Hirasaki M, Nabeshima Y, Nabeshima Y, Katsura Y, et al (2013) In vivo function and evolution of the eutherian-specific pluripotency marker UTF1. PLoS One 8: e68119

Okuda A, Fukushima A, Nishimoto M, Orimo A, Yamagishi T, Nabeshima Y, Kuro-o M, Nabeshima Y, Boon K, Keaveney M, et al (1998) UTF1, a novel transcriptional coactivator expressed in pluripotent embryonic stem cells and extra-embryonic cells. EMBO J 17: 2019–2032

Princz A, Tavernarakis N (2020) SUMOylation in Neurodegenerative Diseases. Gerontology 66: 122–130

Rao S, Zhen S, Roumiantsev S, McDonald LT, Yuan GC, Orkin SH (2010) Differential roles of Sall4 isoforms in embryonic stem cell pluripotency. Mol Cell Biol 30: 5364–5380

Schorova L, Martin S (2016) Sumoylation in Synaptic Function and Dysfunction. Front Synaptic Neurosci 8: 9

Shan SF, Wang LF, Zhai JW, Qin Y, Ouyang HF, Kong YY, Liu J, Wang Y, Xie YH (2008) Modulation of transcriptional corepressor activity of prospero-related homeobox protein (Prox1) by SUMO modification. FEBS Lett 582: 3723–3728

Shevchenko A, Tomas H, Havlis J, Olsen JV, Mann M (2006) In-gel digestion for mass spectrometric characterization of proteins and proteomes. Nat Protoc 1: 2856–2860

Tempe D, Piechaczyk M, Bossis G (2008) SUMO under stress. Biochem Soc Trans 36: 874–878

van den Boom V, Kooistra SM, Boesjes M, Geverts B, Houtsmuller AB, Monzen K, Komuro I, Essers J, Drenth-Diephuis LJ, Eggen BJ (2007) UTF1 is a chromatin-associated protein involved in ES cell differentiation. J Cell Biol 178: 913–924

Yang F, Yao Y, Jiang Y, Lu L, Ma Y, Dai W (2012) Sumoylation is important for stability, subcellular localization, and transcriptional activity of SALL4, an essential stem cell transcription factor. J Biol Chem 287: 38600–38608

Yang Y, He Y, Wang X, Liang Z, He G, Zhang P, Zhu H, Xu N, Liang S (2017) Protein SUMOylation modification and its associations with disease. Open Biol 7:

Young JJ, Kjolby RA, Kong NR, Monica SD, Harland RM (2014) Spalt-like 4 promotes posterior neural fates via repression of pou5f3 family members in Xenopus. Development 141: 1683–1693

Zarelli VE, Dawid IB (2013a) The BTB-containing protein Kctd15 is SUMOylated in vivo. PLoS One 8: e75016

Zarelli VE, Dawid IB (2013b) Inhibition of neural crest formation by Kctd15 involves regulation of transcription factor AP-2. Proc Natl Acad Sci USA 110: 2870–2875

Zhang XD, Goeres J, Zhang H, Yen TJ, Porter AC, Matunis MJ (2008) SUMO-2/3 modification and binding regulate the association of CENP-E with kinetochores and progression through mitosis. Mol Cell 29: 729–741

Zhao Q, Xie Y, Zheng Y, Jiang S, Liu W, Mu W, Liu Z, Zhao Y, Xue Y, Ren J (2014) GPS-SUMO: a tool for the prediction of sumoylation sites and SUMO-interaction motifs. Nucleic Acids Res 42: W325–330

